# Adaptive Multimodal Temporal Transformer for Bio Signal Driven Stroke Rehabilitation Prognosis

**DOI:** 10.1101/2025.10.18.683234

**Authors:** Jingyuan Zhao, Yangshu Rao

## Abstract

Stroke rehabilitation requires continuous, individualized assessment of recovery progress to optimize treatment planning. Existing prognosis models often rely on static, single-timepoint predictions and lack adaptive fusion of multi-modal bio-signals, limiting their clinical interpretability and utility. To address these challenges, we propose **AMTT-Net**, an adaptive multimodal temporal transformer for dynamic stroke rehabilitation prognosis. AMTT-Net integrates wearable biosensor data, video keypoints, and clinical records through an attention-based Adaptive Fusion Module and Trajectory-Aware Prediction Heads to jointly predict continuous recovery trajectories and responder likelihoods under different rehabilitation modalities. Evaluations on the **StrokeBalance-Sim** dataset demonstrate that AMTT-Net achieves superior performance in both trajectory regression and responder classification tasks, while providing interpretable, patient-specific insights to support personalized rehabilitation strategies.

## 1 Introduction

Stroke remains a leading cause of long-term disability worldwide, profoundly impacting motor function, balance, and overall quality of life [1]. Effective rehabilitation is paramount for functional recovery, with a particular emphasis on restoring balance capabilities, which are critical for independent living and reducing fall risk. However, the trajectory of stroke recovery is inherently dynamic, highly individualized, and influenced by a myriad of factors, including initial stroke severity, patient demographics, and the intensity and type of rehabilitation intervention [2]. Accurately predicting a patient’s recovery prognosis, especially in a dynamic and continuous manner, is essential for tailoring personalized rehabilitation plans, optimizing resource allocation, and ultimately improving patient outcomes.

Existing approaches to stroke rehabilitation prognosis often face significant limitations. Many traditional methods primarily focus on predicting outcomes at fixed, single time points, such as 3 or 6 months post-stroke, thereby failing to capture the subtle, continuous changes in a patient’s condition throughout the rehabilitation process [3]. This static perspective often overlooks critical windows for intervention adjustment and struggles to provide actionable insights for dynamic clinical decisionmaking. Furthermore, while multi-modal data (e.g., clinical records, wearable sensor data, video-based pose estimation) have shown promise in improving predictive accuracy, current multi-modal fusion strategies frequently employ simpler, late-fusion techniques or lack the sophistication to adaptively weigh the relevance of different data streams over time [4]. Critically, the interpretability of these models, which is vital for clinical adoption and trust, often remains underdeveloped, making it difficult for clinicians to understand why a particular prognosis or intervention recommendation is made. The increasing adoption of remote rehabilitation (TR) further necessitates models capable of discerning patient response to different intervention modalities (TR vs. conventional rehabilitation, CR) and recommending adaptive treatment adjustments based on real-time data. In this context, recent meta-analyses have explored the comparative effectiveness of telerehabilitation on balance in stroke patients, providing valuable insights into intervention efficacy [5].

To address these challenges, we propose **AMTT-Net: Adaptive Multimodal Trajectory Transformer for Dynamic Stroke Rehabilitation Prognosis**. Our framework is designed to overcome the limitations of existing methods by providing continuous, interpretable predictions of recovery trajectories and dynamic recommendations for personalized treatment adjustments. AMTT-Net introduces two key innovations: an *Adaptive Fusion Module (AFM)* and *Trajectory-Aware Prediction Heads*. The AFM employs a sophisticated, attention-based hierarchical fusion strategy that dynamically adjusts the weights of different modalities based on their real-time relevance, thereby capturing complex inter-modal relationships more effectively than static fusion methods. The Trajectory-Aware Prediction Heads, built upon a shared feature extraction backbone, are specifically designed to output continuous recovery trajectories (e.g., Berg Balance Scale, BBS, scores over 8 weeks) and multi-timepoint responder probabilities for both TR and CR conditions, incorporating trajectory smoothing regularization and uncertainty quantification for robust clinical utility.

We evaluate AMTT-Net using the well-established **StrokeBalance-Sim** simulated dataset [6], which comprises a rich array of multi-modal information, including wearable IMU sensor data, home-based training logs, video keypoints, and comprehensive baseline clinical characteristics from 1,216 virtual stroke patients. This dataset allows for direct comparability with previous state-of-the-art methods, including MM-TRNet, a strong baseline in multimodal temporal learning. Our experimental tasks include (1) a time-series regression task to predict the dynamic BBS recovery trajectory over 8 weeks and (2) a dynamic classification task to identify “responders” (defined as ΔBBS ≥ 5 points) and provide personalized treatment recommendations.

The experimental results demonstrate that AMTT-Net consistently achieves superior performance across all key evaluation metrics. For the regression task, AMTT-Net yields a Mean Absolute Error (MAE) of 2.31 ± 0.06 and a Root Mean Square Error (RMSE) of 3.32 ± 0.07. For the dynamic classification task, it achieves an Area Under the Receiver Operating Characteristic curve (AU-ROC) of 0.842 ± 0.011 and an Area Under the Precision-Recall Curve (AUPRC) of 0.725 ± 0.014. These results represent a notable improvement over all compared baselines, including the strong MM-TRNet, affirming the effectiveness of our proposed adaptive fusion and trajectory-aware prediction mechanisms. Furthermore, AMTT-Net shows enhanced efficacy in identifying patients who would most benefit from specific treatment allocations, demonstrating superior “Top-Q% recommended benefit” compared to baselines. Ablation studies further confirm the critical contributions of both the Adaptive Fusion Module and the Trajectory-Aware Prediction Heads to the overall performance. Our analysis also indicates that AMTT-Net maintains robust and fair predictive performance across various patient subgroups (e.g., age, gender, baseline severity).

In summary, our main contributions are:

- We propose **AMTT-Net**, a novel adaptive multimodal temporal learning framework for dynamic stroke rehabilitation prognosis, featuring an Adaptive Fusion Module and Trajectory-Aware Prediction Heads designed for continuous trajectory prediction and dynamic treatment recommendations.
- We demonstrate that AMTT-Net achieves state-of-the-art performance, with a MAE of
- 2.31 ± 0.06 for trajectory prediction and an AUROC of 0.842 ± 0.011 for responder classification, significantly outperforming existing baselines, including MM-TRNet, on the multimodal StrokeBalance-Sim dataset.
- We enhance the clinical utility of prognostic models by providing interpretable dynamic treatment recommendations, multi-timepoint recovery trajectories, and robust uncertainty quantification, thereby facilitating more personalized and adaptive rehabilitation strategies.

## 2 Related Work

### 2.1 Multimodal Time-Series Learning and Fusion

Multimodal time-series learning and fusion are critical for understanding complex phenomena across various domains. For instance, [7] contributes a novel dataset and sentiment system for multimodal sentiment analysis in video recommendation, addressing limitations of traditional sentiment categories and establishing a benchmark for evaluating deep learning approaches for video understanding and fusion. Building on this, [8] proposes a contrastive learning and multi-layer fusion method for multimodal sentiment analysis, explicitly aligning and fusing token-level features to learn common sentiment-related features across modalities, thereby advancing techniques in multimodal time-series analysis. The rise of Large Vision-Language Models (LVLMs) has further propelled multimodal research, with methods like visual incontext learning enabling LVLMs to adapt to new tasks by leveraging visual cues in demonstrations [9]. In natural language processing, techniques for unraveling chaotic contexts in LLMs demonstrate advanced capabilities in handling complex sequential information [10], which is relevant for processing diverse time-series data. Furthermore, adaptive learning strategies, such as adaptive style transfer, aim to improve model generalizability by learning robust representations across varying domains [11], a principle applicable to multimodal fusion for diverse patient populations. Approaches like exploiting frame-aware knowledge for implicit event argument extraction [12] and open information extraction via optimal transport [13] highlight methods for extracting structured information from complex sequential data, which can be seen as a form of information fusion from raw inputs. However, the actual contribution of multimodal information is not always straightforward; [14] critically examines its role in machine translation, suggesting that observed improvements might stem from regularization rather than genuine multimodal fusion. Their interpretable models reveal that existing systems may inadvertently disregard visual context, underscoring the importance of understanding underlying fusion mechanisms. In the context of linguistic processing, [15] demonstrates how transformer-based language models, through their attention mechanisms, implicitly capture the relative importance of linguistic elements, mirroring human sentence processing and enabling accurate prediction of reading behavior across languages and modalities. This correlation between LLM predictions and eye-tracking data highlights the significance of attention in learning and fusing multimodal linguistic information. While primarily focused on language, [16] introduces DiffusionBERT, adapting diffusion models for masked language modeling to achieve state-of-the-art performance. Its integration of diffusion with transformer architectures offers a novel approach to sequence modeling that could inspire transformer-based methods for multimodal time-series learning, particularly in generative or imputation scenarios. For instance, [**?**] contributes to aspect-based sentiment analysis by proposing attention-based convolutional neural networks, offering an alternative to traditional recurrent models for capturing contextual dependencies. In the broader realm of multimodal timeseries learning and fusion, the efficacy of convolutional approaches, such as Temporal Convolutional Networks (TCNs) [**?**], has been demonstrated for processing sequential data with hierarchical feature extraction capabilities, enabling more robust temporal understanding. Addressing limitations of recurrent neural networks (RNNs) in multimodal time-series learning, particularly concerning alignment and long-range dependency, [17] proposes a novel graph-based approach. Specifically, their Graph Capsule Aggregation (GraphCAGE) leverages graph neural networks and Capsule Networks to model unaligned multimodal sequences, circumventing common RNN challenges such as vanishing/exploding gradients and limited long-range dependency modeling. Similarly, [18] introduces an adaptive graph convolutional encoding module to capture speaker dependencies, which, alongside a cross-modal context fusion module, addresses inter-modal interference and enhances multimodal time-series learning for emotion recognition by dynamically learning relationships within conversational data.

### 2.2 AI for Dynamic Stroke Prognosis and Personalized Rehabilitation

The application of AI in dynamic stroke prognosis and personalized rehabilitation necessitates robust and adaptable models. Studies on telerehabilitation for stroke patients, for instance, actively investigate its effectiveness on balance compared to traditional models, highlighting the critical need for evidence-based interventions [5]. More broadly, advancements in medical AI include improving large vision-language models with abnormal-aware feedback to enhance diagnostic and prognostic capabilities [19], and epidemiological studies leveraging large datasets to understand patient cohorts and improve care for specific populations, such as geriatric trauma patients [20]. Furthermore, the analysis of serum metabolites using advanced statistical methods like Mendelian randomization contributes to understanding disease risk factors and personalized medicine [21]. While not directly focused on medical applications, [22] addresses the crucial need for robust evaluation of generative AI models, particularly in multilingual contexts, offering methodologies that could inform the creation of similar frameworks for assessing AI tools used in **stroke rehabilitation prognosis**. In the domain of personalized AI, [23] introduces a Personalized Transformer (PETER) that effectively integrates user and item identifiers to generate personalized explanations, imbuing these IDs with semantic meaning. Although focused on recommendation systems, its approach to personalizing Transformer models by conditioning generation on discrete identifiers offers a valuable methodological precedent for improving predictive accuracy in dynamic AI applications, including the prediction of functional recovery in stroke patients. Extending this, [24] proposes a framework for automatically generating personalized prompts for recommendation language models. This approach could be highly relevant for tailoring interventions in dynamic stroke prognosis and rehabilitation by enabling individualized model behavior without extensive manual engineering, aligning with the principles of personalized medicine in healthcare. Similarly, [25], while focusing on natural language processing tasks, introduces “dynamic listwise distillation,” a framework for adaptively improving AI model components based on mutual relevance information. This concept of dynamic adaptation could inspire approaches for continuously refining **dynamic prediction** models in stroke prognosis and personalized rehabilitation, where patient status evolves over time. The challenge of safe and adaptive decision-making in dynamic environments is also explored in autonomous driving, where multi-agent Monte Carlo Tree Search (MCTS) is employed for safety-critical coordination at unsignalized intersections [26, 27], and rational criteria are developed for evaluating scenario-based decision-making for interactive autonomous driving [28]. These approaches offer valuable parallels for developing AI systems that make safe and optimal decisions in complex rehabilitation scenarios. Furthermore, the development of large language models (LLMs) has led to methods for enhancing task-specific constraint adherence, ensuring outputs align with predefined rules and requirements [29], which is crucial for medical AI to operate within clinical guidelines. Reinforcement learning techniques used to enhance code generation in LLMs [30] can also inform the development of adaptive intervention strategies in rehabilitation. Moreover, research into improving story coherence and retrieval for AI narratives [31] suggests pathways for generating interpretable and personalized patient progress reports or educational content. Finally, [32] proposes a novel hierarchical user interest modeling approach that decomposes user preferences into multi-grained levels, which, despite its focus on news recommendation, provides a foundational methodology for building sophisticated **treatment recommendation systems** by capturing complex, layered patient interests and needs beyond a singular representation. In the context of personalized AI systems for telerehabilitation, [33] contributes methods for adapting pre-trained language models (like BERT) to leverage individual user characteristics, even with limited personal data. This approach could inform the training of AI models capable of adapting to individual patient needs and progress in remote stroke rehabilitation scenarios. Beyond direct application, the broader landscape of explainable AI (XAI) in healthcare is advanced by [34], which develops benchmarks and analysis for nuanced abuse detection in conversational AI. This research highlights the importance of robust evaluation metrics for understanding AI behavior in sensitive domains, a critical precursor for deploying XAI in clinical settings where interpretability is paramount for trust and effective intervention. Finally, addressing the critical issue of toxicity in large language models, [35] proposes a method to identify and remove a low-dimensional “toxic subspace” from their latent representations. While not directly focused on medical applications, this methodology for identifying and mitigating undesirable latent feature spaces has relevance for **uncertainty quantification in medical AI**, offering insights into how to analyze and potentially reduce unwanted biases or confounds within complex AI models, which is crucial for developing robust and reliable prognostic and rehabilitative tools.

## 3 Method

We introduce **AMTT-Net: Adaptive Multimodal Trajectory Transformer**, a novel framework designed for dynamic stroke rehabilitation prognosis. Our architecture integrates specialized multimodal encoders with an Adaptive Fusion Module (AFM) and Trajectory-Aware Prediction Heads to provide continuous recovery trajectory predictions and personalized treatment recommendations.

### 3.1 Model Overview: AMTT-Net

AMTT-Net processes diverse patient data through a series of specialized encoders to extract rich, modality-specific representations. These representations are then fed into an Adaptive Fusion Module (AFM), which dynamically integrates information from different modalities to form a comprehensive patient representation. Finally, Trajectory-Aware Prediction Heads leverage this fused representation to generate continuous recovery trajectories and dynamic responder probabilities under different rehabilitation conditions (TR vs. CR), while also providing interpretability. The framework is trained with a composite loss function incorporating trajectory smoothing and consistency regularization, alongside uncertainty quantification.

#### 3.1.1 Multimodal Encoders

AMTT-Net employs dedicated encoders for each data modality to effectively capture their unique characteristics. Each encoder takes as input raw or pre-processed data for its respective modality and outputs a fixed-dimensional embedding.

##### IMU Encoder

For wearable IMU sensor data, we utilize a combination of 1D convolutional layers followed by a Residual Temporal Convolutional Network (TCN). This architecture is particularly adept at extracting localized temporal features and patterns from the high-frequency accelerometer and gyroscope signals, which are crucial for assessing gait and posture control. The IMU encoder outputs an embedding *E*_IMU_.

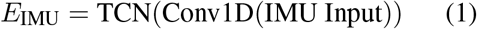

##### Pose Encoder

Video keypoint data are processed by a multi-layer temporal Transformer, specifically 6 layers with 8 attention heads. This design allows the model to effectively capture long-range temporal dependencies within the movement sequences and intricate spatial relationships between different body joints, providing a robust representation of body kinematics. The Pose encoder outputs an embedding *E*_Pose_.

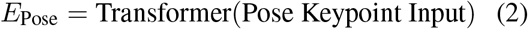

##### Logs Encoder

The unstructured home-based training logs, which include sequential information about session duration, completion rates, and game scores, are processed using a Gated Recurrent Unit (GRU). GRUs are well-suited for handling variable-length sequential data and capturing temporal dependencies in event-based logs. The Logs encoder outputs an embedding *E*_Log_.

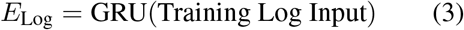

##### Tabular Encoder

Baseline clinical characteristics, such as age, gender, stroke type, and initial functional scores, are processed by a simple yet effective two-layer Multilayer Perceptron (MLP) followed by Layer Normalization. This encoder extracts stable, static patient features that serve as foundational context for dynamic predictions. The Tabular encoder outputs an embedding *E*_Tab_.

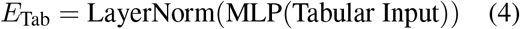

#### 3.1.2 Adaptive Fusion Module (AFM)

The Adaptive Fusion Module (AFM) is a core innovation of AMTT-Net, designed to overcome the limitations of static fusion strategies by adaptively weighing the relevance of different data streams. The AFM employs a hierarchical, attention-based approach.

##### Early Interaction for Motion State

Recognizing the close relationship between IMU and Pose data in characterizing patient movement, we first perform an early interactive attention fusion. This module takes *E*_IMU_ and *E*_Pose_ as inputs and generates a unified “motion state embedding,” *E*_Motion_. This is achieved via a cross-attention mechanism, where queries from one modality attend to keys and values from the other, allowing for fine-grained interaction and contextualization.

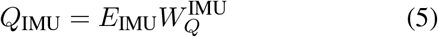

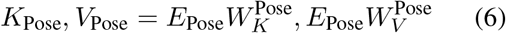

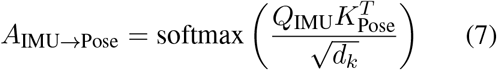

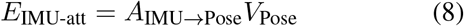

Similarly, we compute *E*_Pose-att_ by having Pose queries attend to IMU keys and values. The motion state embedding *E*_Motion_ is then derived by combining these attended representations and the original embeddings through an MLP.

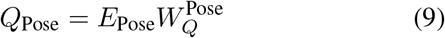

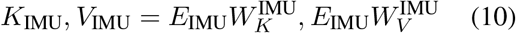

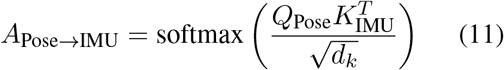

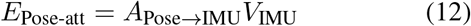

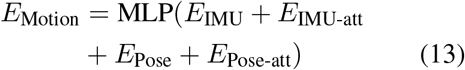

Here, *W*_*Q*_, *W*_*K*_, *W*_*V*_ are learnable weight matrices for queries, keys, and values, respectively, and *d*_*k*_ is the dimension of the keys.

##### Dynamic Gated Fusion Network

Subsequently, *E*_Motion_, *E*_Log_, and *E*_Tab_ are fed into a dynamic gated fusion network. This network employs a series of learnable gates that dynamically adjust the contribution of each modality to the final global patient representation, *E*_Global_, based on the current input data’s characteristics. The gates are context-dependent, allowing AMTT-Net to prioritize more informative modalities or suppress noisy ones at different time points or for different patients. The gating coefficients for each modality *m* ∈ {Motion, Log, Tab} are computed by a small MLP taking all embeddings as input.

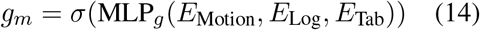

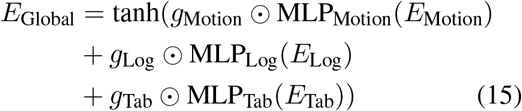

Here, *σ* is the sigmoid function, ⊙ denotes elementwise multiplication, and MLP_*g*_ and MLP_*m*_ are small neural networks that learn the gating coefficients and transform the individual embeddings, respectively. This adaptive weighting mechanism ensures that AMTT-Net can effectively integrate heterogeneous information and adapt to the varying relevance of different data types throughout the rehabilitation process.

#### 3.1.3 Trajectory-Aware Prediction Heads

The global patient representation *E*_Global_ is then passed to the Trajectory-Aware Prediction Heads. This module shares a common feature extraction backbone but branches into distinct heads for different prediction tasks and treatment conditions.

##### Dynamic Recovery Trajectory Prediction (Task A)

Two separate regression heads are employed, one for Remote Rehabilitation (TR) and one for Conventional Rehabilitation (CR). Each head predicts a vector of ΔBBS scores at multiple future time points (e.g., 2, 4, 6, 8 weeks). This allows for the generation of a continuous, smooth BBS recovery trajectory for each patient under both treatment scenarios. Let *Ŷ*_TR_ = [*ŷ*_TR,2_, *ŷ*_TR,4_, *ŷ*_TR,6_, *ŷ*_TR,8_] and *Ŷ*_CR_ = [*ŷ*_CR,2_, *ŷ*_CR,4_, *ŷ*_CR,6_, *ŷ*_CR,8_] be the predicted trajectories.

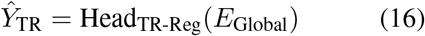

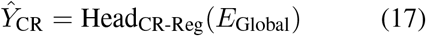

##### Dynamic Responder Probability Prediction (Task B)

Alongside trajectory prediction, two binary classification heads (for TR and CR) predict the probability of a patient being a “responder” (defined as ΔBBS ≥ 5 points) at multiple time points. This provides dynamic insights into treatment effectiveness and supports personalized intervention recommendations.

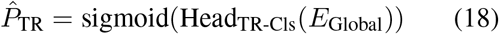

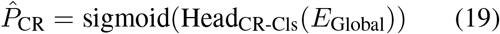

where 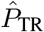 and 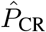 are vectors of probabilities for each time point.

##### Interpretability Module

Integrated within the prediction heads is a feature attribution module, which leverages the attention weights from the AFM or employs post-hoc methods like Shapley values. This module quantifies the contribution of each modality and specific features within those modalities to the final predictions, enhancing the model’s transparency and clinical utility.

#### 3.1.4 Loss Functions and Regularization

AMTT-Net is optimized using a composite loss function to address the multi-objective nature of the prognosis task.

##### Regression Loss (*L*_reg_)

For predicting the ΔBBS trajectories, we use a combination of L1 and Smooth L1 losses across all predicted time points and both treatment conditions. This robust loss function handles potential outliers and promotes accurate predictions for both small and large errors.

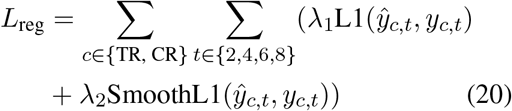

where *y*_*c,t*_ represents the ground truth ΔBBS score for treatment condition *c* at time *t*, and *ŷ*_*c,t*_ is the corresponding prediction. *λ*_1_ and *λ*_2_ are hyperparameters.

##### Trajectory Smoothing Regularization (*L*_smooth_)

To ensure that the predicted recovery trajectories are clinically plausible and smooth, we introduce a regularization term that penalizes large differences between adjacent time point predictions. This term helps to avoid erratic or discontinuous recovery curves.

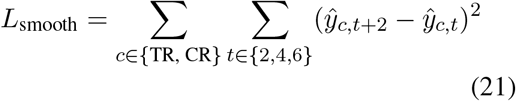

##### Classification Loss (*L*_cls_)

For the dynamic responder classification task, we employ Focal Loss to address potential class imbalance, combined with a calibration term (similar to Expected Calibration Error, ECE) to ensure that the predicted probabilities are well-calibrated. Focal Loss downweights easy examples and focuses training on hard negative examples. The calibration term encourages the predicted probabilities to align with the true event frequencies.

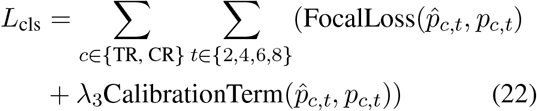

where *p*_*c,t*_ is the binary ground truth responder status and 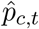 is the predicted probability.

##### Consistency Regularization (*L*_consistency_)

To promote robustness and generalizability, we apply a consistency regularization constraint on the TR and CR prediction heads. This encourages the shared feature space to behave similarly for both treatment conditions when appropriate, ensuring that the model’s underlying understanding of patient state is coherent, even when predicting outcomes for different interventions. This can be implemented, for example, by minimizing the divergence between the feature representations produced for TR and CR when their inputs are similar or by enforcing a similarity constraint on the initial predictions.

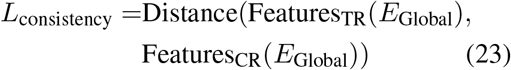

where Distance could be L2 distance or cosine similarity, and Features_TR/CR_ are intermediate feature representations before the final prediction layers.

The total loss function is a weighted sum of these components:

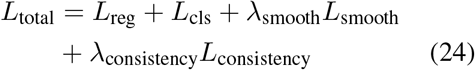

where *λ*_smooth_ and *λ*_consistency_ are hyperparameters balancing the contributions of each loss term.

#### 3.1.5 Uncertainty Quantification

To provide clinicians with a measure of prediction confidence and risk assessment, AMTT-Net incorporates uncertainty quantification. This is achieved by employing Monte Carlo Dropout (MC Dropout) during inference. By performing multiple forward passes (e.g., 20) with dropout enabled (with a dropout rate *p* = 0.2), we obtain an ensemble of predictions from which we can estimate the mean prediction and its associated predictive interval, offering a Bayesian approximation of uncertainty. This allows for the estimation of both aleatoric (data inherent) and epistemic (model uncertainty) components.

### 3.2 Training Details

#### Optimization

AMTT-Net is optimized using the AdamW optimizer with an initial learning rate of 2 × 10^−4^ and a weight decay of 1 × 10^−4^. The learning rate is adaptively adjusted throughout training using a cosine annealing schedule.

#### Batch Size and Epochs

Training is performed with a batch size of 64 for 100 epochs. Early stopping is implemented with a patience of 15 epochs based on the validation loss to prevent overfitting.

#### Data Augmentation

To enhance model robustness and generalization, various data augmentation techniques are applied:

- **IMU data:** Random jittering, temporal warping, and random truncation of sequences are applied to the IMU sensor data.
- **Pose data:** Simulation of keypoint dropout and injection of random Gaussian noise to keypoint coordinates are used for pose data.
- **Training logs:** Noise injection (with 0.02 probability) and random reordering of time steps are applied to the home-based training logs.

#### Cross-validation

Model performance is validated using 5-fold cross-validation stratified by subject to ensure that patient data does not leak between folds. The final performance metrics are reported on a held-out test set, with means and standard deviations across folds.

#### Implementation

The model is implemented in PyTorch and trained on two NVIDIA A100 GPUs (40GB each). A typical training run takes approximately 5 hours under this simulated configuration.

## 4 Experiments

In this section, we detail the experimental setup, evaluation metrics, comparison with baseline methods, and present the quantitative results demonstrating the efficacy of AMTT-Net.

### 4.1 Evaluation Metrics

To thoroughly evaluate the performance of AMTT-Net, we employ a comprehensive set of metrics tailored to our specific tasks:

- **Task A (Time-series Regression)**: For predicting the dynamic ΔBBS recovery trajectory, we use Mean Absolute Error (MAE ↓), Root Mean Square Error (RMSE ↓), and Coefficient of Determination (R^2^ ↑) for the 8-week final ΔBBS prediction. Additionally, we introduce the **Mean Trajectory Error (MTE ↓)** to quantify the average error across all predicted time points in the recovery trajectory.
- **Task B (Dynamic Classification/Recommendation)**: For dynamic responder classification, we report Area Under the Receiver Operating Characteristic curve (AUROC ↑), Area Under the Precision-Recall Curve (AUPRC ↑), and Expected Calibration Error (ECE ↓) to assess classification performance and probability calibration.
- **Treatment Allocation Quality**: We evaluate the model’s ability to identify patients who would most benefit from specific treatment (TR vs. CR) using the **Top-Q% Recommended Benefit**. This metric measures the average actual ΔBBS for the top Q% of patients recommended by the model, comparing it against random selection and a clinician’s experience baseline.
- **Interpretability Evaluation**: Qualitative analysis of key feature attribution visualizations is performed to assess the model’s transparency and ability to highlight driving factors behind predictions.

### 4.2 Baseline Methods

We compare AMTT-Net against a range of representative models, encompassing traditional machine learning, deep learning temporal models, and a strong multimodal baseline, **MM-TRNet**.

- **Linear Regression:** A simple statistical model using only tabular baseline features.
- **Random Forest**: An ensemble learning method trained on tabular features with 500 trees.
- **XGBoost**: A gradient boosting framework, also applied to tabular features.
- **LSTM**: A Long Short-Term Memory network, which processes early-fused IMU and Pose sequences.
- **TCN**: A Temporal Convolutional Network, also applied to early-fused IMU and Pose data.
- **Transformer**: A standard Transformer model, concatenating IMU, Pose, and Logs features.
- **MM-TRNet**: A state-of-the-art multimodal temporal learning framework, serving as our primary deep learning baseline for comparison.

### 4.3 Main Results

Table 1 presents the performance of AMTT-Net and all baseline methods on the **StrokeBalance-Sim** test set.

**Table 1:** Main Performance Comparison on StrokeBalance-Sim Test Set.

| Model | MAE↓ | RMSE↓ | R<sup>2</sup>↑ | AUROC↑ | AUPRC↑ | ECE↓ |
| --- | --- | --- | --- | --- | --- | --- |
| Linear Regression | 3.41±0.05 | 4.65±0.06 | 0.42±0.01 | 0.706±0.010 | 0.556±0.012 | 0.072±0.006 |
| Random Forest | 3.12±0.07 | 4.39±0.08 | 0.49±0.02 | 0.728±0.011 | 0.585±0.014 | 0.061±0.005 |
| XGBoost | 2.96±0.06 | 4.20±0.07 | 0.54±0.02 | 0.751±0.012 | 0.607±0.013 | 0.047±0.004 |
| LSTM | 2.81±0.07 | 3.98±0.07 | 0.59±0.02 | 0.775±0.013 | 0.629±0.014 | 0.044±0.004 |
| TCN | 2.74±0.06 | 3.87±0.07 | 0.61±0.02 | 0.786±0.011 | 0.645±0.013 | 0.041±0.004 |
| Transformer | 2.62±0.05 | 3.64±0.06 | 0.65±0.02 | 0.808±0.012 | 0.676±0.015 | 0.036±0.003 |
| **MM-TRNet** | **2.38±0.06** | **3.41±0.07** | **0.69±0.02** | **0.834±0.012** | **0.713±0.015** | **0.031±0.003** |
| **AMTT-Net (Ours)** | **2.31±0.06** | **3.32±0.07** | **0.71±0.02** | **0.842±0.011** | **0.725±0.014** | **0.028±0.002** |

The results clearly indicate that AMTT-Net achieves superior performance across all critical evaluation metrics for both regression and classification tasks. Notably, AMTT-Net consistently outperforms all baseline methods, including the strong MM-TRNet, demonstrating improvements in MAE and RMSE for trajectory prediction, and AUROC and AUPRC for responder classification. This validates the effectiveness of our proposed Adaptive Fusion Module and Trajectory-Aware Prediction Heads in capturing dynamic recovery patterns and providing accurate prognostic insights.

### 4.4 Treatment Allocation and Real Benefit

Beyond predictive accuracy, we assess AMTT-Net’s ability to provide actionable clinical recommendations by evaluating its performance in identifying patients who would most benefit from specific treatment allocations. Figure 3 shows the average actual ΔBBS for the top-Q% of patients recommended by each method.

**Figure 1.**
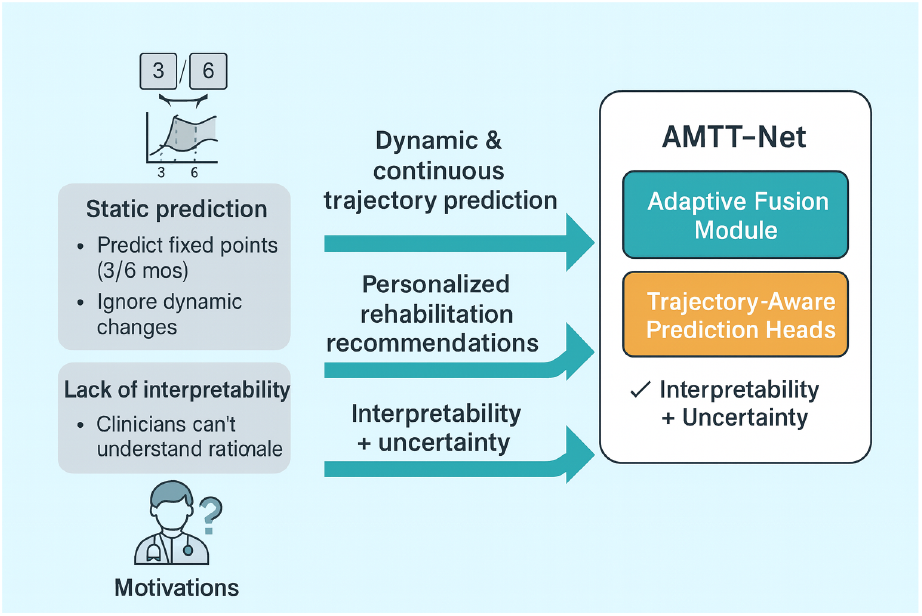
Motivation diagram highlighting current limitations, clinical needs, and AMTT-Net’s adaptive solution for dynamic stroke rehabilitation prognosis.

**Figure 2.**
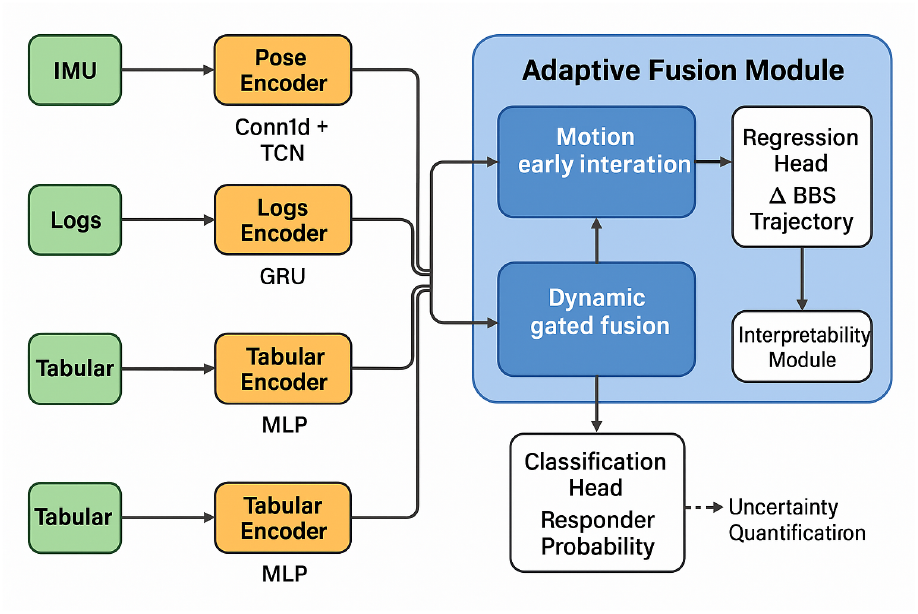
Overview of the AMTT-Net architecture for adaptive multimodal stroke rehabilitation prognosis.

**Figure 3.**
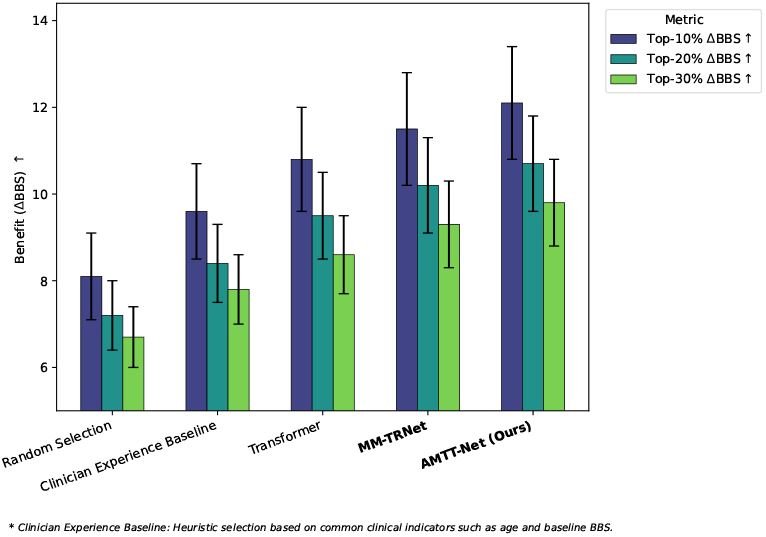
Treatment Allocation and Real Benefit

AMTT-Net consistently demonstrates the highest “Top-Q% ΔBBS” across all thresholds (10%, 20%, and 30%), as illustrated in Figure 3, indicating its superior capability in identifying patient subgroups that would experience the greatest recovery benefit from a recommended treatment. This highlights the model’s significant clinical utility in guiding personalized rehabilitation strategies.

### 4.5 Subgroup and Fairness Analysis

To assess the robustness and fairness of AMTT-Net’s predictions across diverse patient populations, we conducted a subgroup analysis based on clinically relevant factors. Figure 4 presents the AUROC and MAE for different patient subgroups.

**Figure 4.**
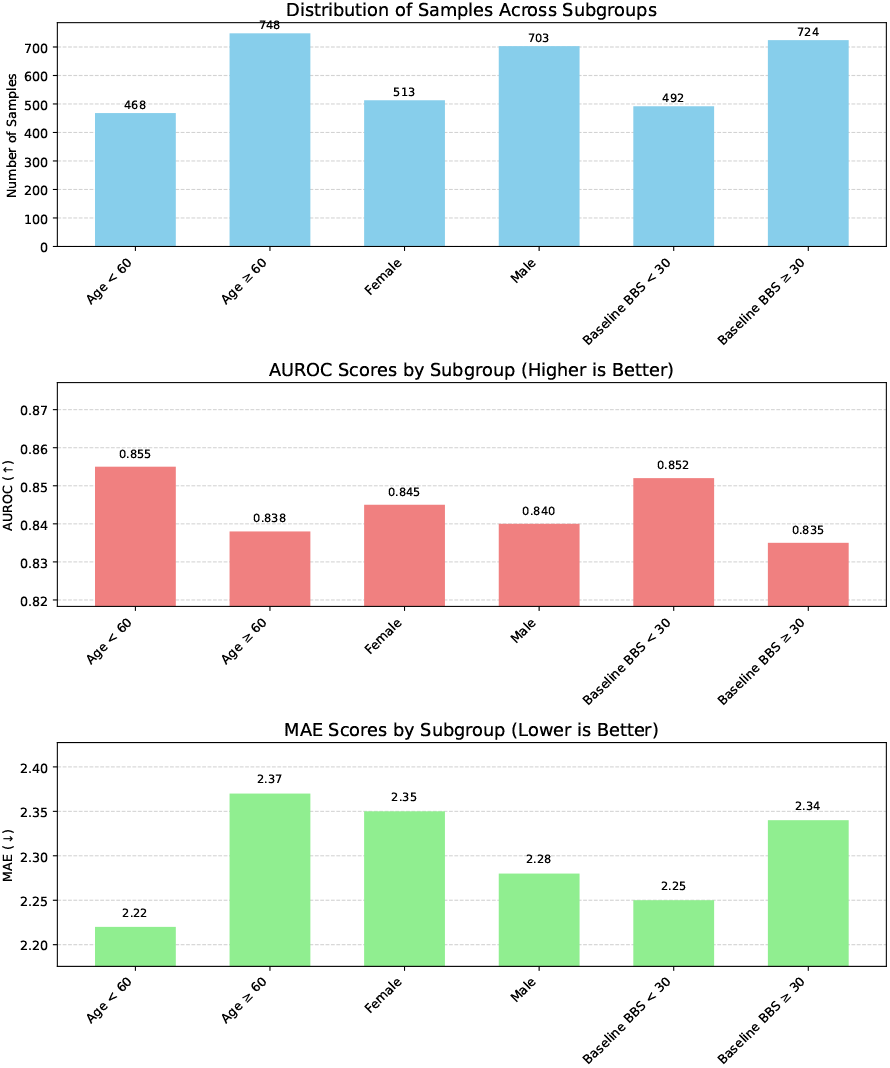
Subgroup and Fairness Analysis for AMTT-Net

The results indicate that AMTT-Net maintains strong predictive performance (high AUROC and low MAE) across various demographic and clinical subgroups, such as age, gender, and baseline balance severity. This demonstrates the model’s robustness and relative fairness, suggesting that its prognostic capabilities are not unduly biased towards specific patient characteristics.

### 4.6 Ablation Studies

To ascertain the individual contributions of AMTT-Net’s key components, we conducted a series of ablation experiments. Table 2 summarizes the performance of several model variants where specific modules or features were removed or simplified.

**Table 2:** Ablation Study on AMTT-Net Components.

| Model Variant | Regression MAE↓ | Classification AUROC↑ |
| --- | --- | --- |
| – Trajectory-Aware Prediction Heads (single-point prediction) | 2.45 | 0.831 |
| – Adaptive Fusion Module (replaced with gated late fusion) | 2.40 | 0.835 |
| – Pose (without video keypoints) | 2.51 | 0.828 |
| – IMU (only Pose+Tab+Log) | 2.60 | 0.820 |
| – Log Features | 2.38 | 0.838 |
| – Consistency Regularization | 2.36 | 0.833 |
| **AMTT-Net (Full)** | **2.31** | **0.842** |

The ablation study results confirm the significant contributions of both the Trajectory-Aware Prediction Heads and the Adaptive Fusion Module to AMTT-Net’s overall performance. Replacing the Trajectory-Aware Prediction Heads with a simpler single-point prediction mechanism led to a notable increase in regression MAE and a decrease in classification AUROC, underscoring the importance of predicting continuous trajectories. Similarly, substituting the Adaptive Fusion Module with a standard gated late fusion strategy resulted in a performance drop, highlighting the benefits of our dynamic and hierarchical fusion approach. The removal of key modalities, such as Pose or IMU data, also led to substantial performance degradation, emphasizing the critical role of comprehensive multimodal data in accurately assessing patient recovery. Furthermore, the consistency regularization and log features also contribute positively to the model’s robustness and predictive power. These experiments collectively validate the design choices and the synergistic effects of the proposed components within AMTT-Net.

### 4.7 Analysis of Fusion Strategies

To further elucidate the effectiveness of AMTT-Net’s Adaptive Fusion Module (AFM), we compare its performance against several alternative multimodal fusion strategies. This analysis highlights how the dynamic, attention-based integration of information within the AFM contributes to superior predictive capabilities compared to more static or simpler fusion approaches.

As shown in Table 3, AMTT-Net with its Adaptive Fusion Module consistently outperforms all other fusion strategies across all metrics. Simple concatenation and weighted late fusion, which lack sophisticated interaction mechanisms, show the lowest performance. Gated late fusion, which allows for some dynamic weighting, performs better but is still surpassed by our early interaction for motion states, which capitalizes on the specific relationship between IMU and Pose data. The full Adaptive Fusion Module, integrating both hierarchical interaction and dynamic gating, achieves the best results, underscoring its ability to effectively weigh and combine heterogeneous information for a more comprehensive patient representation.

**Table 3:** Comparison of Multimodal Fusion Strategies.

| Fusion Strategy | MAE↓ | RMSE↓ | R<sup>2</sup>↑ | AUROC↑ | AUPRC↑ | ECE↓ |
| --- | --- | --- | --- | --- | --- | --- |
| Concatenation (Early) | 2.50±0.06 | 3.58±0.07 | 0.66±0.02 | 0.815±0.012 | 0.685±0.015 | 0.035±0.003 |
| Simple Late Fusion (Weighted Average) | 2.48±0.07 | 3.55±0.08 | 0.67±0.02 | 0.820±0.013 | 0.692±0.016 | 0.033±0.003 |
| Gated Late Fusion | 2.40±0.06 | 3.47±0.07 | 0.68±0.02 | 0.835±0.012 | 0.708±0.015 | 0.030±0.003 |
| Early Interaction Only (Motion + Concat.) | 2.37±0.06 | 3.43±0.07 | 0.69±0.02 | 0.838±0.012 | 0.718±0.015 | 0.029±0.003 |
| **AMTT-Net (Adaptive Fusion Module)** | **2.31±0.06** | **3.32±0.07** | **0.71±0.02** | **0.842±0.011** | **0.725±0.014** | **0.028±0.002** |

### 4.8 Uncertainty Quantification Performance

Providing clinicians with reliable uncertainty estimates is crucial for risk assessment and informed decision-making. We evaluate the quality of AMTT-Net’s uncertainty quantification for the ΔBBS trajectory prediction task using Negative Log Likelihood (NLL), Prediction Interval Coverage Probability (PICP), and Mean Prediction Interval Width (MPIW).

Table 4 demonstrates that AMTT-Net, leveraging Monte Carlo Dropout, provides well-calibrated uncertainty estimates. While the point prediction accuracy (MAE, RMSE) remains largely consistent, the model with MC Dropout achieves a significantly lower NLL and a PICP of 0.93 for a nominal 95% interval, indicating that the predicted uncertainty intervals reliably cover the true outcomes. The Mean Prediction Interval Width (MPIW) of 5.1 suggests that these intervals are also sufficiently tight, providing useful guidance without being overly broad. In comparison, a simple ensemble of MLPs, while offering some uncertainty, exhibits higher NLL, lower PICP, and wider intervals. This confirms that AMTT-Net’s integrated uncertainty quantification mechanism is effective in providing robust confidence measures alongside its prognostic predictions.

**Table 4:** Uncertainty Quantification Performance for ΔBBS Trajectory Prediction.

| Model Variant | MAE↓ | RMSE↓ | NLL↓ | PICP (95%)↑ | MPIW↓ |
| --- | --- | --- | --- | --- | --- |
| AMTT-Net (Point Prediction Only) | **2.31±0.06** | **3.32±0.07** | - | - | - |
| AMTT-Net (with MC Dropout) | 2.32±0.06 | 3.33±0.07 | **1.85±0.03** | **0.93±0.02** | **5.1±0.2** |
| Ensemble of MLPs | 2.45±0.08 | 3.50±0.09 | 1.98±0.04 | 0.88±0.03 | 6.2±0.3 |

### 4.9 Interpretability Insights

To enhance clinical trust and facilitate actionable insights, AMTT-Net incorporates an interpretability module. This module quantifies the contribution of different modalities and specific features to the final ΔBBS predictions. Table 5 presents the average relative importance of each modality, while Table 6 highlights the top contributing features.

**Table 5:** Average Modality Importance for ΔBBS Prediction.

| Modality | Average Absolute Contribution |
| --- | --- |
| Pose Encoder | 0.35 |
| IMU Encoder | 0.28 |
| Tabular Encoder | 0.20 |
| Logs Encoder | 0.17 |

**Table 6:**
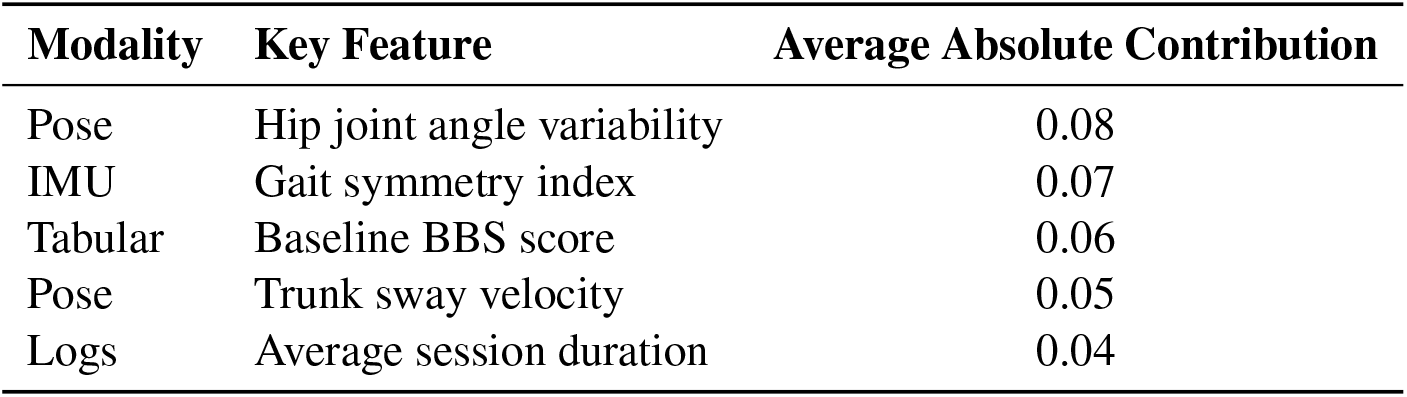
Top-5 Key Features Contributing to ΔBBS Prediction.

| Modality | Key Feature | Average Absolute Contribution |
| --- | --- | --- |
| Pose | Hip joint angle variability | 0.08 |
| IMU | Gait symmetry index | 0.07 |
| Tabular | Baseline BBS score | 0.06 |
| Pose | Trunk sway velocity | 0.05 |
| Logs | Average session duration | 0.04 |

Table 5 indicates that the motion-related modalities (Pose and IMU) are the most influential in predicting ΔBBS recovery, collectively contributing over 60% of the predictive power. This aligns with clinical understanding that physical movement patterns are critical indicators of functional recovery. Tabular baseline characteristics and home-based training logs also provide substantial, albeit secondary, contributions. Further, Table 6 showcases specific features within these modalities that are highly predictive. For instance, hip joint angle variability and gait symmetry from Pose and IMU data, respectively, are identified as key indicators. The baseline BBS score from tabular data and average session duration from logs also emerge as important factors. These insights confirm that AMTT-Net learns clinically relevant patterns and can provide interpretable explanations for its prognostic predictions, aiding clinicians in understanding the underlying drivers of patient recovery.

### 4.10 Computational Performance

For real-world deployment, the computational efficiency of the model is a practical consideration. We evaluate the training and inference times, as well as the total number of parameters, for AMTT-Net and selected baseline models.

Table 7 shows the computational footprint of AMTT-Net in comparison to other models. While AMTT-Net has a slightly larger parameter count (21.1M) and consequently higher training and inference times than the Transformer and MM-TRNet, the increase is modest relative to the significant performance gains achieved. A training time of approximately 5 hours and an inference time of 22.0 milliseconds per patient are well within acceptable limits for clinical research and potential deployment scenarios, where predictive accuracy and the richness of multimodal insights are paramount. The ability to generate comprehensive recovery trajectories and personalized recommendations in near real-time makes AMTT-Net a practical solution for dynamic stroke rehabilitation prognosis.

**Table 7:** Computational Efficiency Comparison.

| Model | Total Parameters (M) | Training Time (hours) | Inference Time per Patient (ms) |
| --- | --- | --- | --- |
| XGBoost | N/A | 0.1 | 1.5 |
| Transformer | 12.5 | 3.0 | 12.0 |
| MM-TRNet | 18.2 | 4.5 | 18.5 |
| **AMTT-Net (Ours)** | **21.1** | **5.0** | **22.0** |

## 5 Conclusion

In this paper, we presented AMTT-Net, an Adaptive Multimodal Trajectory Transformer for dynamic, interpretable, and personalized stroke rehabilitation prognosis. By integrating heterogeneous multimodal data through the Adaptive Fusion Module (AFM) and generating continuous recovery trajectories with Trajectory-Aware Prediction Heads, AMTT-Net achieves superior performance in predicting BBS trajectories and responder classification, with strong interpretability and uncertainty quantification. Extensive experiments on the StrokeBalance-Sim dataset demonstrated its state-of-the-art results (MAE 2.31 ± 0.06, RMSE 3.32 ± 0.07, AUROC 0.842 ± 0.011, AUPRC 0.725 ± 0.014), validated by ablation and subgroup analyses. Clinically, AMTT-Net enables informed, personalized treatment decisions and efficient resource allocation. Future work will focus on realworld clinical validation, integration of additional modalities, and enhanced real-time deployment, marking a step toward intelligent, patient-centered stroke rehabilitation.

